# Why does a flying fish taxi on sea surface before take-off? A hydrodynamic interpretation

**DOI:** 10.1101/765560

**Authors:** Jian Deng, Shuhong Wang, Lingxin Zhang, Xuerui Mao

## Abstract

Flying fish have been observed jumping out of warm ocean waters worldwide. Before take-off, the flying fish are seen to taxi on the water surface by rapidly beating their semi-submerged tail fins, which process may help them airborne with enough speed to glide over a long distance. To understand the underlying physical mechanisms, here, we study a flying fish, 0.25 m in length and 0.191 kg in weight, considering both its underwater swimming and surface taxiing locomotion. Its hydrodynamic characteristics are numerically studied by computational fluid dynamics (CFD). Underwater, the fish is assumed to swim at a constant speed of 10 m s^−1^. Different critical frequencies are identified for various maximum deflected angles, ranging from *θ*_0_ = 10° to 30°, at which the fish reaches cruising states, when the horizontal forces are balanced. The corresponding minimum power required for cruising swimming is 350 W, obtained at a deflected angle of 10° and a critical frequency of 145 Hz. In contrast, in the taxiing stage, the minimum power required for a stead-state locomotion at 10 m s^−1^ is 36 W, occurring at a deflected angle of 15° and a frequency of 50 Hz. We note that the power is significantly smaller than the swimming locomotion. Further, by increasing the flapping power, we find that larger speeds can be achieved. In specific, when the power is brought up to 350 W, it can reach a speed of 16.5 m s^−1^. Clearly, from the direct comparison between the two locomotive modes, it is apparently evidenced that the flying fish can be further accelerated by taxiing along the water surface.

## I. INTRODUCTION

In recent years, there is a rising interest in new concepts of robots with hybrid and multi-modal locomotion (Low et al. 2015), learning from their natural counterparts. For example, the so-called amphibians, such as turtles and salamanders can swim in water and walk on land, while swans can swim on water and fly effectively. More interestingly, there is a family of marine fish, exocoetidae, in the order Beloniformes, class Actinopterygii, known colloquially as flying fish (Breder Jr 1938, Nelson et al. 2016). They do not fly, in the sense of flapping their wing-sized pectoral fins, but actually perform unpowered glide. Their remarkable abilities could inspire innovative designs to improve the way man-made systems operating in the environments consisting of multiple media. However, it is very challenging to design such aerial-aquatic robots or Aquatic Micro Aerial Vehicles (AquaMAVs), mimicking real flying fish. The challenges lie in platform design, high power density propulsion systems and control across the air-water interface, due to the impossibility to simultaneously optimize the performance in different locomotive modes (Gao and Techet 2011). It is therefore necessary to identify the key design principles that make their mobility realizable and effective by understanding the underlying physics of multi-modal locomotion (Low et al. 2015).

According to the statistic data provided by zoologists, adult flying fish vary in size from the two-winged Parexocoetus brachypterus with a maximum recorded standard length (*L*) of 125 mm to the four-winged Cypsilurus lineatus with a maximum length of 378 mm (Bruun 1935). It was reported that an adult four-winged flying fish, 0.3 m in length, can reach a speed of up to 10 m s^−1^ (about 20-30 body lengths s^−1^) in water with its pectoral fins folded tightly against its streamlined body (Davenport 1994). Upon piercing the sea surface, it spreads its enlarged pectoral fins and gains additional thrust by beating its tail fins with the lower lobes submerged at a frequency of 50-70 strokes s^−1^. When sufficiently high speed has been attained, the tail is lifted clear of the water and the fish is airborne, gliding a few meters above the surface at a peak speed of about 15 – 20 m s^−1^ (Davenport 1994). Field observations have revealed that a flying fish can reach a gliding distance of 50 m and a peak height up to 8 m, when the tail fin is held high and still (Hubbs 1933). The flying fish can make several consecutive glides, with its tail propelling it up again each time it sinks back to the surface. A total flight distance of 400 m can be achieved in 30 s by the repeated taxiing before the fish submerges eventually into the water (Franzisket 1965).

Flying fish were thought to have evolved the remarkable gliding ability to escape predators (Davenport 1994), which is intuitively attractive since the predator may lose sight of the flying fish when it bursts into air. However, some scientists believed that the periodic flights of flying fish could also be part of an energy-saving strategy akin to some marine mammals which repeatedly jump out of the water when cruising for long distance, yielding additional benefits of being a more efficient mode of transportation (Rayner 1986). It has been reported that animals use diverse strategies to reduce the energetically expensive cost of locomotion, ranging from morphological to behavioural solutions (Schmidt-Nielsen 1972). Intermittent locomotion, as a specific behavioural strategy is widely taken by both vertebrates and invertebrates to reduce the cost of movement (Kramer and McLaughlin 2001). For example, the intermittent locomotion, analogous to undulating flight, of a bird involves gliding with flexed wings interspersed with active flapping, in which the potential energy from gravity and altitude is translated into horizonal distance via gliding, resulting in savings of mechanical power compared with continuous level flight (Gleiss et al. 2011). Similarly, the aerial-aquatic locomotion of a flying fish can also be regarded as a strategy for power energy saving, whilst in a more unique way. It is apparent that flying fish are likely to experience less resistance in air than that in water assuming moving forward at the same speed. Since the flying stage is unpowered, the flying fish should, first, be equipped with highly modified pectoral fins providing sufficient lift during gliding, which has been proved by previous experiments (Park and Choi 2010) as well as our numerical simulations (Deng et al. 2019). Second, a large take-off speed is preferred for the flying fish to achieve the horizontal distance of gliding flight as long as possible. To achieve the second goal, taxiing on the water surface seems to be a reasonable choice, which as we have introduced above, can probably accelerate the flying fish to a speed up to 15 – 20 m s^−1^ (Davenport 1994).

From the point of view of physiology, the swimming ability of a flying fish 0.3 m in length swimming at a maximum cruising speed of 10 m s^−1^, is extraordinary. Wardle (1975) found that the maximum ‘steady-state’, or cruising speed of fish could be predicted empirically by

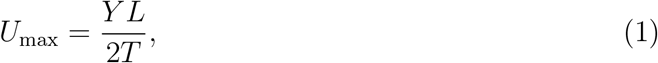

where *Y* is the stride length (the ratio between an one-tail-beating forward distance and the body length *L*), and *T* is the time for one contraction of the swimming muscles (two contraction time is equal to one tail beating cycle). *Y* varies from 0.6 to 0.81. Therefore, for the flying fish of *L* = 0.3 m with a stride length of 0.8, and cruising at 10 m s^−1^:

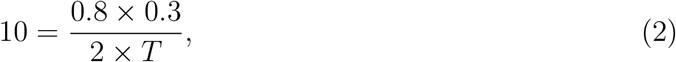

then *T* = 0.012 s is obtained, which means that the flying fish has to perform 83.3 contractions s^−1^ or 41.7 tail beats s^−1^. Unfortunately, according to Wardle’s curve (Wardle 1975) for the relationship between *L* and *T* for various ambient temperatures, the minimum contraction time is about 0.025 s for *L* = 0.3 m (at 20 °*C*). Therefore, the required *T* value is about half the predicted from Wardle’s curve (Wardle 1975). It is thereby unlikely that flying fish will be able to emerge and fly at temperatures lower than 20 °*C*. Indeed, the tail beating rates of about 50 beats s^−1^ have been recorded in warmer waters (in Caribbean waters at about 25 °*C*), indicating that the contraction rate is within the capability of flying fish (Hertel 1966).

Hydrodynamically, as an approximation, we can evaluate the drag force, then the power required to achieve a ‘steady-state’ swim by using the dead-drag coefficient of a fish of similar size, i.e., *C*_*D*_ ≈ 0.02 (at *Re* = 2.54 × 10^6^) (Blake 1983). The drag force *F*_*D*_ and the corresponding power *P*_*r*_ can be expressed as

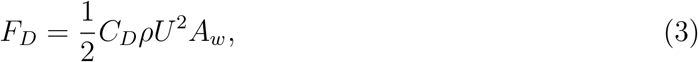

and

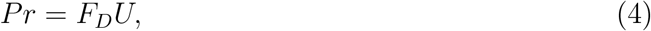

respectively. For a flying fish with *L* = 0.25 m, swimming at *U* =10 m s^−1^, excluding the pectoral fins (which will be folded against the body during swimming), assuming the wing area *A*_*w*_ = 0.023 m^2^ and *ρ* = 1000 kg m^−3^ (Gao and Techet 2011), we get *F*_*D*_ = 23 N and *Pr* = 230 W. As suggested by Gao and Techet (2011), this required power for the flying fish, weighing 0.2 kg, relates to a muscle power density of 2300 W kg^−1^ (assuming 50% muscle by weight). We note that this power density is far beyond the range of the existing artificial actuators, from electromagnetic actuators such as DC motors (on the order of 100 W kg^−1^) to pneumatic actuators and air muscles (weight ratios of up to 400 W kg^−1^).

It appears that the maximum swimming speed of 10 m s^−1^ for a flying fish is markedly high from the perspectives of hydrodynamic resistance and power requirement. It is unlikely that the flying fish will be able to swim faster, due to the quadratical increase of the drag force with the swimming speed. We therefore conjecture that taxiing on the water surface is a necessary stage for the flying fish to reach an ideal take-off speed, though more solid evidence is required, which is the central aim of the current study. Moreover, despite the great challenges associated with the design of aerial-aquatic robots, there is limited understanding of the underlying physical mechanisms for their natural counterparts.

In this paper, we study a flying fish with both underwater swimming and water-surface taxiing locomotion considered. We aim to explain why a flying fish taxis on the water surface before take-off from the perspective of hydrodynamics, with a particular concern on direct comparisons in beating frequency and power requirement between the two locomotive modes.

## II. PHYSICAL MODEL AND NUMERICAL METHOD

### A. Physical model

We consider two different locomotive modes of a flying fish, i.e., underwater swimming and water surface taxiing, as shown in figure 1. The pectoral fins are folded for the swimming mode (see figure 1(a) and (b)), while spread for the taxiing mode (see figure 1(c)). For both modes, the pectoral fins are with straight leading edge, following the same geometry used in our previous study (Deng et al. 2019). As a simplification, the fins are very thin with simple rectangular cross sections, with a thickness of 1 mm, which can thereby be neglected in the analysis, and the pelvic fins are removed.

**FIG. 1:**
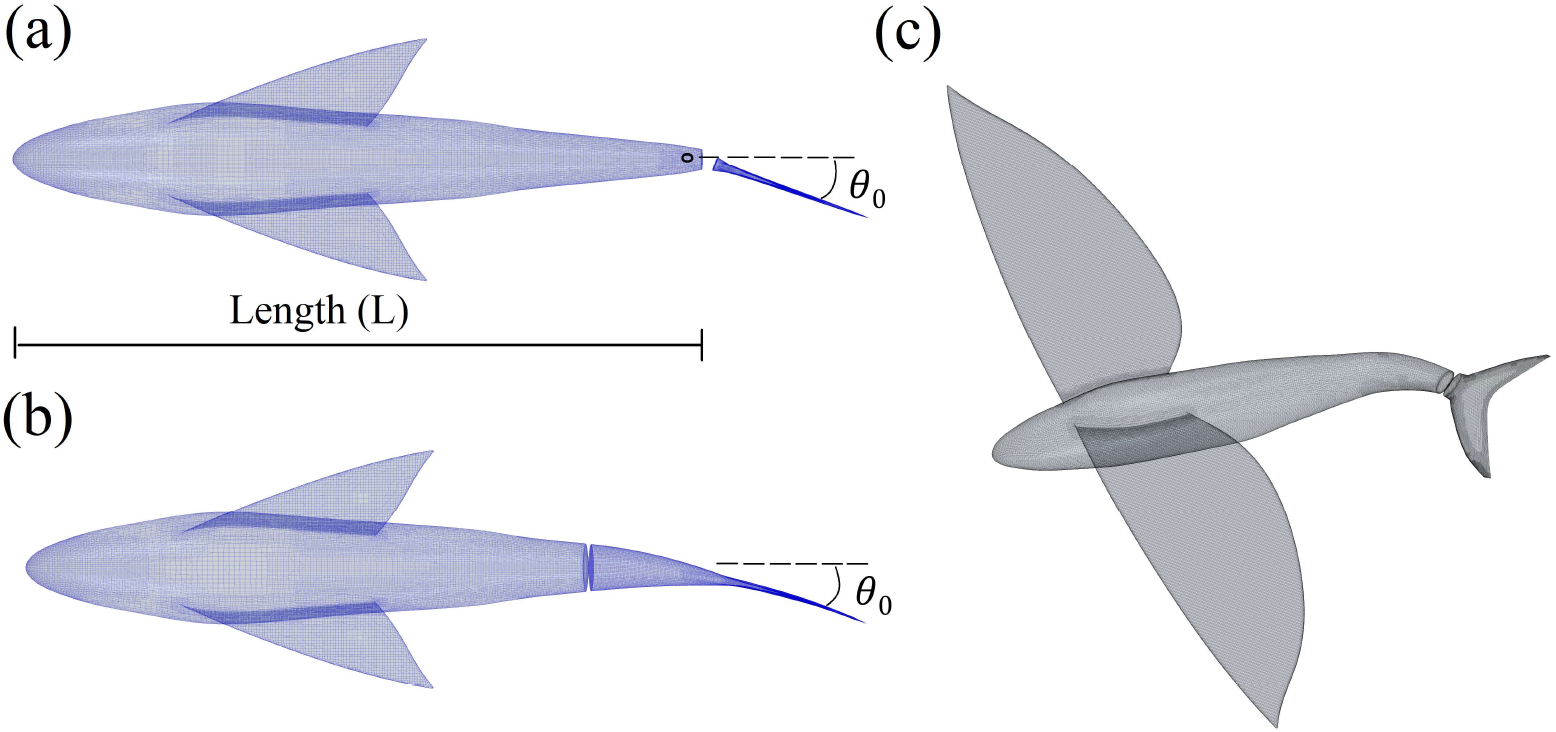
Geometrical configurations of the three flying fish models: underwater swimming models with (a) a rigid flapping tail and (b) a periodically morphing tail, and (c) the taxiing model with a spreading pectoral fins. Note that (a) and (b) are shown at the time instant of 1/4*T* when the tails reach their maximum deflected angles (*θ*_0_ = 20°), where *T* is the period of one beating cycle.

The morphological parameters are chosen as follows: the standard length *L* = 0.25 m, the wing area for the spread pectoral fins *A* = 0.024 m^2^, the wing span *S* = 0.47 m, the wing aspect ratio *AR* = 9.2 (*AR* = *S*^2^/*A*), the average wing chord length *C* = 0.051 m. The body mass is set to *W* = 0.191 kg, resulting in a wing loading of 78 N m^−2^, falling in the range of wing loadings for six genera reported by Fish (1990). The root chord length of the fish tail is *D* = 0.03 m, measuring from the pivoting point to the tail fork point. We note that the currently adopted flying fish model follows closely the model B that we chose previously for a gliding flight (Deng et al. 2019), while differing from the model B in standard length, which was *L* = 0.2 m.

For the rigid flapping tail, it pitches periodically along a pivoting point as marked in figure 1(a), while for the periodically morphing tail, it deforms laterally and periodically with the amplitude of lateral displacement increasing linearly from the pivoting point to the tail tips, as shown in figure 1(b). A maximum deflected angle *θ*_0_ is defined to quantize the beating amplitude of the tail.

The angles of attack (*α*) are 0° and 5° for the swimming and taxiing locomotion, respectively. For the taxiing locomotion as shown in figure 1(c), the tail is bent down to an inclined angle of 30° with respect to the horizontal plane, therefore the lower lobe of the tail fin is submerged in the initial flow field.

### B. Numerical method

The numerical simulations are carried out using the commercial CFD code STAR-CCM+ 12.06.011 (CD-adapco 2017), which is based on the finite volume method. The governing equations for the incompressible, viscous flow include a continuity equation and momentum equation for each of the three dimensions. A segregated flow model is used to solve each of the momentum equations in turn, one for each dimension. The linkage between the momentum and continuity equations is achieved with a predictor-corrector approach. A hybrid second-order upwind/bounded-central scheme is used for the convection term, with the upwind blending factor set to 0.15. For temporal discretization, a first-order implicit scheme is used, with several inner iterations involved in each physical time step to converge the solution for the given time step.

Volume of fluid (VOF) method is used to model the free surface for the surface taxxing cases. The high-Resolution Interface Capturing (HRIC) scheme is applied to the convective terms of the VOF transport equation, resulting in a scheme that is suited for sharp interface tracking (Muzaferija and Peric 1999). To model the turbulence, the SST K-Omega Detached Eddy model is used, which combines the features of SST K-Omega RANS model in the boundary layers with a large eddy simulation (LES) in unsteady separated regions (Shur et al. 2008).

To deal with the flapping or periodically morphing tail, an overset meshing technique is adopted (Hadzic 2006). The computational domain includes a background mesh enclosing the entire solution domain, containing the pectoral fins and the fish body, and one smaller overset mesh (a cubic box) containing the tail fin, as shown in figure 2. We note that the mesh shown in figure 2 is for the surface taxiing cases, while the swimming cases follow the same strategy of mesh generation, except for the less requirement of cell number due to the folded pectoral fins. For the cases of rigid flapping tail, the entire overset mesh moves with the tail, while for the cases of morphing tail, the vertices in the overset mesh redistribute in response to the movement of the fish tail. The cells in the background mesh covered by the overset region are deactivated and do not take part in the simulations. These cells can be anyway reactivated later on, if by means of movements they become active again. At the boundaries between the background and overset regions, active (discretization) and interpolation cells are present. The solution is computed at the active cells, while it is interpolated at the interpolation cells. Detailed implementation of the overset techniques can be found in Ref.(Hadzic 2006) or the STAR-CCM+ manual (CD-adapco 2017).

**FIG. 2:**
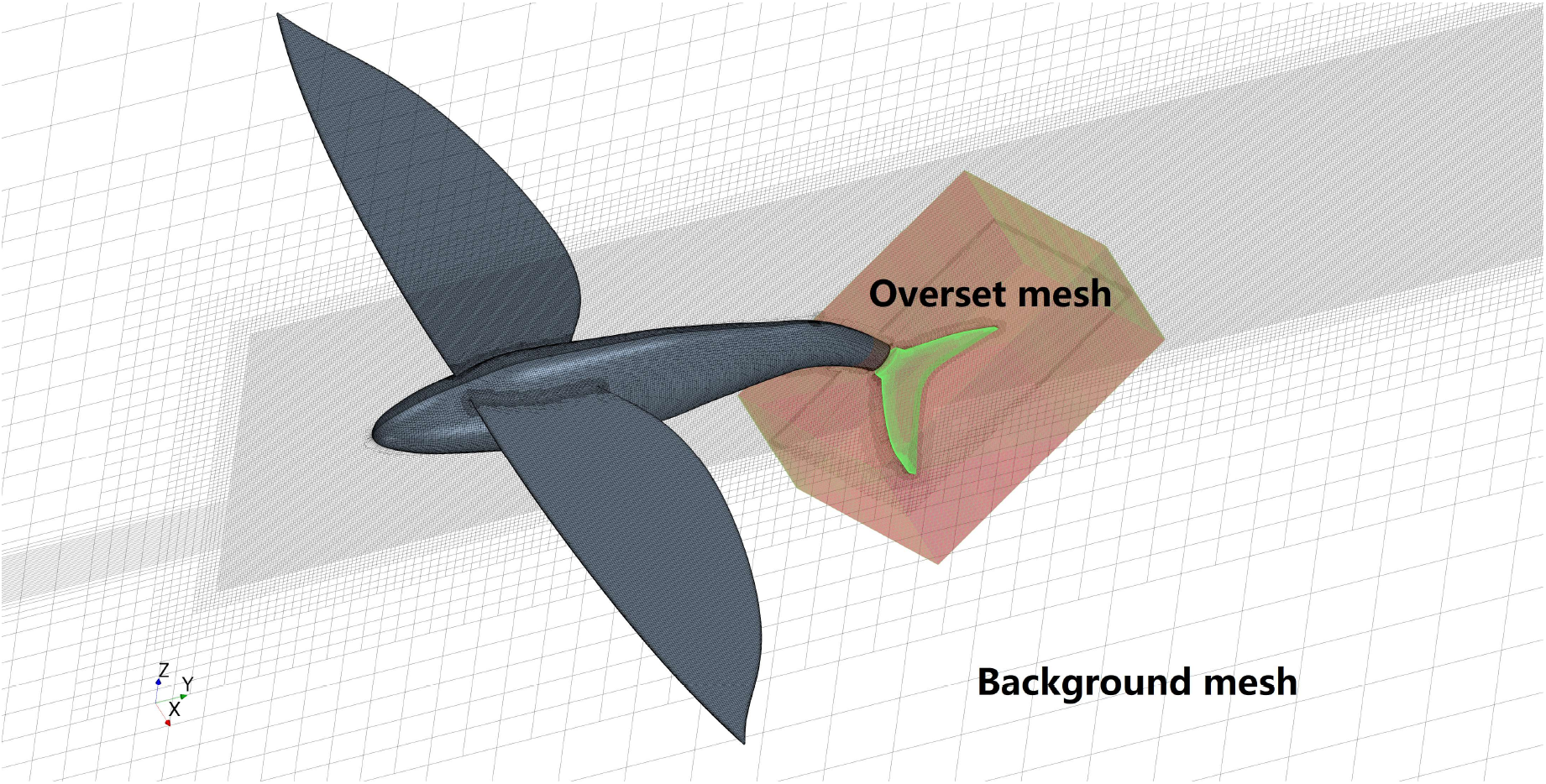
Computational domain showing the background mesh and overset mesh, and the mesh refinement around the flying fish as well as that along the free surface.

### C. Computational setup

The computational domain is a cuboid, of which the dimensions of its outer boundary is 2 m × 4 m × 2 m at the x-y-z coordinates, as shown in figure 2, for both the swimming and taxiing locomotion. In the streamwise, or y direction, inlet and outlet boundaries are 1 m and 3 m respectively away from the fish nose. The grid cells on the surfaces, particularly around the fins, of the flying fish, and along the free surface are refined, as shown in figure 2. Five layers of cells are generated within the wall boundaries, with the first layer with the height of about 0.0003 m.

The velocity magnitude at the inlet boundary is set to a constant value, for example *U*_∞_=10 m s^−1^. The density and dynamic viscosity for water are *ρ*_1_ = 997.56 kg m^−3^ and *μ*_1_ = 8.887×10^−4^ kg m^−1^ s^−1^, respectively. They are *ρ*_2_ = 1.18 kg m^−3^ and *μ*_2_ = 1.855×10^−5^ kg m^−1^ s^−1^ respectively for air. Therefore, the resulting Reynolds number in the water *Re*_1_ = *ρ*_1_*U*_∞_*L*/*μ*_1_ = 2.8 × 10^6^, taking the standard length as its length scale, and that in the air *Re*_2_ = *ρ*_2_*U*_∞_*C*/*μ*_2_ = 3.2 × 10^4^, taking the chord length as its length scale, which is a relatively low Reynolds number in contrast to artificial aircrafts.

### D. Validation

To resolve the unsteadiness of the flow induced by flapping tail, first, we set the time step size to be 1/(1000*f*), i.e., 1000 time steps per flapping cycle, which yields time-accurate predictions for both mean and instantaneous values. Moreover, the time step size Δ*t* is adjusted during simulations to meet the Courant-Friedrichs-Lewy (CFL) condition, i.e., *Co* = Δ*t*|**U**|/Δ*x* < 1. To obtain mesh independence results, we choose a typical surface taxiing case to perform rigorous self-consistency tests. Different mesh resolutions are used, from a coarse background mesh (6, 522, 198 cells) to a very fine mesh (63, 834, 110 cells), with the overset mesh adjusted accordingly. In this case, the flying fish taxis at *U*_∞_ = 10 m s^−1^ and *θ*_0_ = 15°, and the tails beats at a frequency of *f* = 100 Hz. In table I, we show the time-averaged thrust force *F*_*T*_ = −*F*_*y*_ and the power input *P*_*in*_, which is calculated by integrating the inner product of distributed forces and moving velocities along the tail surface. The relative errors are evaluated with respect to the finest mesh resolution. It can be seen that our medium-mesh resolution provides satisfactory accuracy in space, as far as thrust force and power input concerned. Indeed, for our test cases, differences between the medium-mesh (33, 267, 959 cells) and the fine-mesh (63, 834, 110 cells) results are quite small, with variations of less than 1% on both thrust force and power input. Therefore, we use the medium mesh of 33, 267, 959 cells for all the following calculations of taxiing locomotion. For the underwater swimming cases, since the pectoral fins are folded (see figure 1(a) and (b)), requiring less mesh requirement around them, and there is no free surface to perform mesh refinement, the corresponding mesh numbers are greatly reduced, which are 9, 777, 998 for the background mesh and 981, 935 for the overset mesh, by adopting the same mesh generation strategy.

**TABLE I:** Results of validation through space refinement; taxiing at *U*_∞_ = 10 m s^−1^, *θ*_0_ = 15° and *f* = 100 Hz

| Cell number for<br>background mesh | Cell number for<br>overset mesh | Thrust force<br>($F_T$)(N) | Power input<br>($P_{in}$)(W) | Error for<br>$F_T$ | Error for<br>$P_{in}$ |
| --- | --- | --- | --- | --- | --- |
| 6,522,198 | 476,650 | 7.39 | 362 | 6.5% | 5.8% |
| 9,448,796 | 584,526 | 7.28 | 355 | 4.9% | 3.8% |
| 25,465,732 | 885,864 | 7.03 | 349 | 1.3% | 2.0% |
| 33,267,959 | 1,006,804 | 6.96 | 340 | 0.3% | -0.5% |
| 49,123,779 | 1,556,511 | 6.93 | 344 | 0.1% | 0.6% |
| 63,834,110 | 1,998,974 | 6.94 | 342 | 0% | 0% |

## III. RESULTS AND DISCUSSION

### A. Underwater swimming locomotion

For the swimming locomotion, the pectoral fins are folded, as shown in figure 1 (a) and (b). Two propulsive modes are considered, both swimming at a constant speed of *U*_∞_ = 10 m s^−1^. First, the tail performs a periodically pitching motion (the rigid flapping mode) with a sinusoidal variation of the deflected angle with time. For the second one, the morphing mode, the grid points on the tail oscillate laterally (in the x direction) and periodically, with the amplitude of oscillation increasing linearly from the pivoting point to the tail tips, mimicking the undulating motion of a fish tail. Since we have defined a maximum deflected angle *θ*_0_ (see figure 1 (a) and (b)), which can be regarded as a characteristic width for the wake dynamics, it is possible to make a direct comparison between these two propulsive modes.

Figure 3(a) shows the variations of thrust forces with beating frequency for the rigid flapping mode, with four different *θ*_0_ considered. It is seen that the thrust force increases along with the beating frequency for all cases. We find that, for each case, the thrust force crosses the zero value line at a specific critical frequency, indicating the transition from a deceleration state to an acceleration swimming state. At these critical points, the horizontal forces are balanced, therefore the flying fish reaches a steady-state or cruising locomotion. We should point out that the thrust force *F*_*T*_ considered here is calculated by integrating the distributed pressure and viscous forces along all parts, including the fish body, folded pectoral fins and the tail. It is observed that the critical frequencies for transition are around 115 Hz for *θ*_0_ = 15°, 20° and 30°. However, for the small deflected angle, *θ*_0_ = 10°, the transition occurs at a markedly higher frequency of *f* = 145 Hz. It suggests that the fish should beat its tail at a higher frequency if the amplitude has been reduced. We can define a non-dimensional frequency, or the Strouhal number as

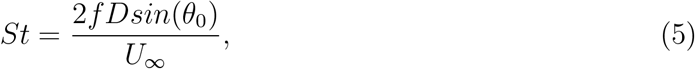

in which *D* =0.03 m (defined in section II A), then the deceleration-to-acceleration transitions occur in the range of *St* = 0.151 to 0.345. This range accords surprisingly well with the cruise Strouhal numbers, lying within a narrow interval 0.2 < *St* < 0.4, for a wide range of flying and swimming animals (Taylor et al. 2003). It is noted that our previously identified Strouhal number, *St* = 0.225, of a pure pitching foil for drag-thrust transition also lies in these ranges (Deng et al. 2015, 2016).

**FIG. 3:**
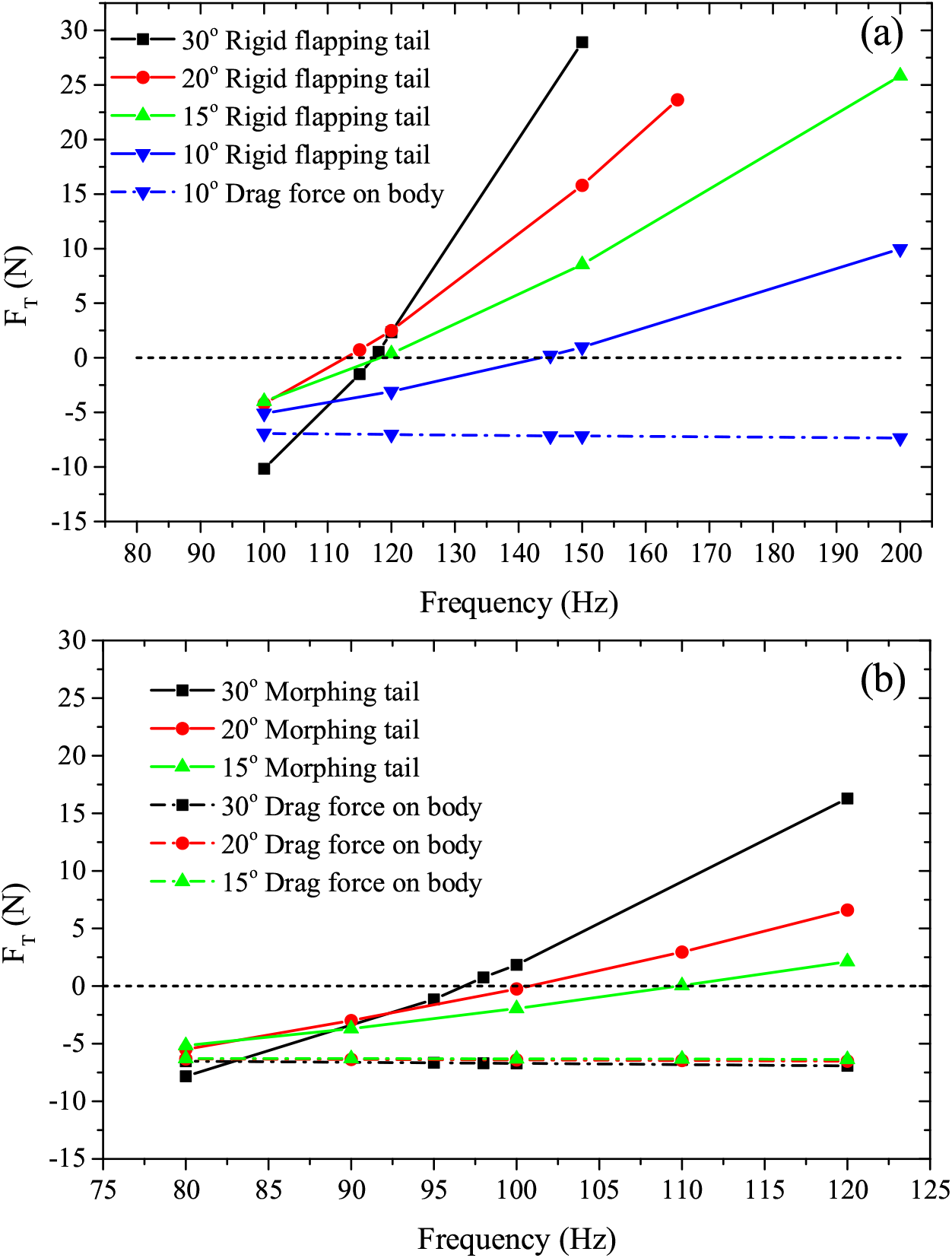
Thrust forces on the flying fish when it swims at a constant speed of 10 m s^−1^ for different beating frequencies, propelled by (a) rigid flapping tail; (b) periodically morphing tail.

For the periodically morphing tail, the critical frequencies are 110 Hz, 100 Hz and 98 Hz respectively for *θ* = 15°, 20° and 30°, or in the range of Strouhal number *St* = 0.152 to 0.33, as shown in figure 3(b). It appears that the propulsive mode does not change significantly the locomotive performance, which is mainly determined by the wake width. In figure 4, we show the wake topologies represented by Q iso-surfaces. The Q-criterion (Jeong and Hussain 1995) defines a vortex as a spatial region where

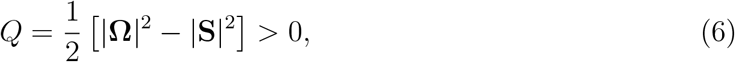

where 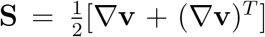 is the rate of strain tensor, and 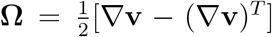 is the vorticity tensor. The positive value of *Q* means that the Euclidean norm of the vorticity tensor dominates that of the rate of strain. In figure 4, we highlight the vortex cores using iso-surfaces of *Q* = 7.5(*U*_∞_/*L*)^2^. Positive values of *Q* give prominence to regions of high swirl in comparison to shear to represent coherent vortices. The wake width is mainly determined by the flapping amplitude, as clearly seen in In figure 4. In figure 4 (a), when the deflected angle is small, the vortical structures are confined to a very narrow lateral space, while for the larger deflected angles, the flow wakes are wider, as seen in figure 4 (b) and (c). Unlike traditional Bénard-von Kármán (BvK) vortex streets viewed in a two-dimensional plane (Deng and Caulfield 2015), here the flow wakes produced by the forked tail (caudal fin) show strongly three-dimensionality. Nevertheless, the reversed BvK streets signaling propulsive wakes can still by observed. As shown noticeably in figure 4 (b) and (c), the vortices formed from one side of the tail fin shed to the other side of the wake, forming two rows of vortex streets, which are connected by vortex filaments in the braid region.

**FIG. 4:**
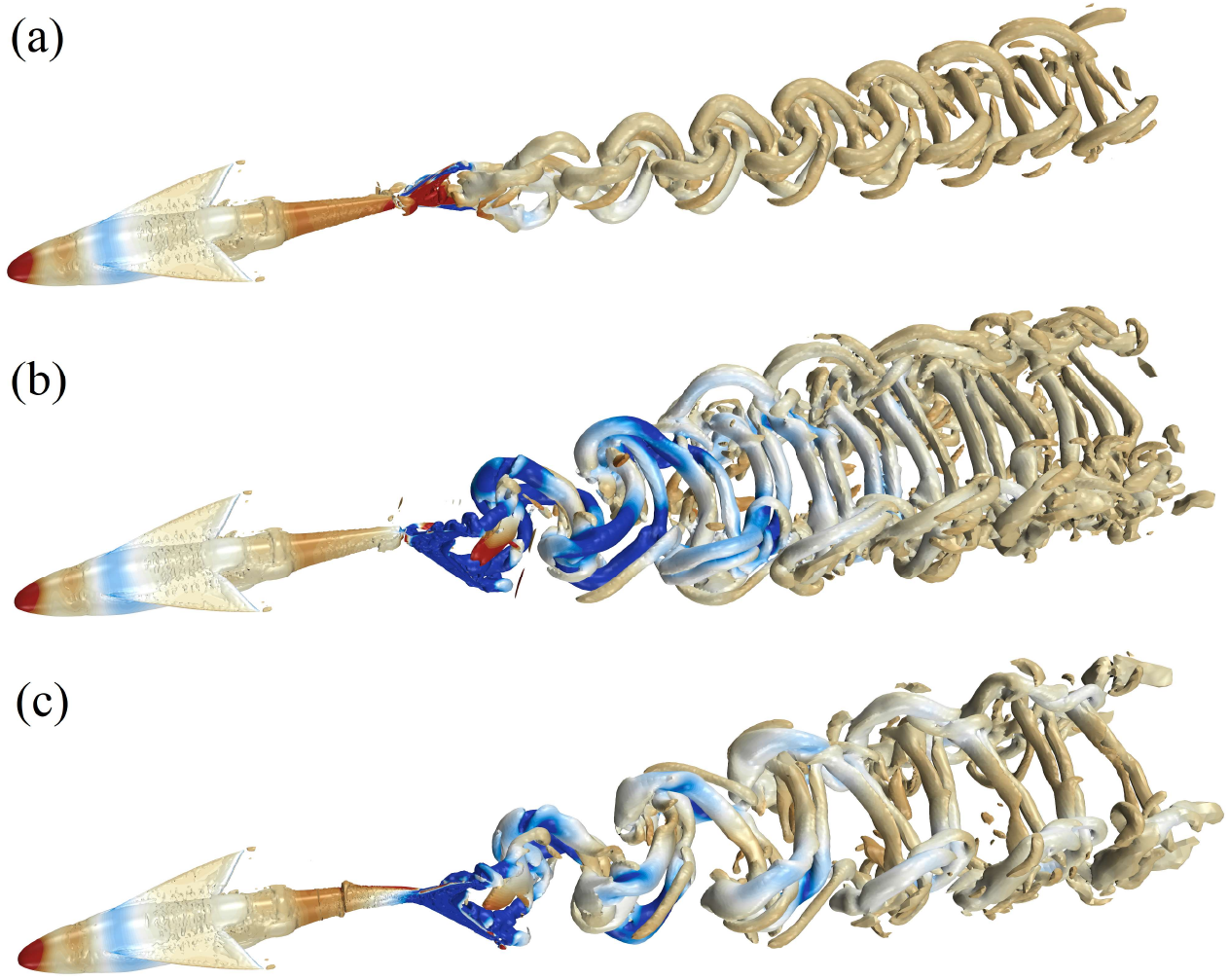
Wake topologies for the underwater swimming flying fish, visualized using *Q* = 7.5(*U*_∞_/*L*)^2^, in which *U*_∞_ =10 m s^−1^ is the velocity magnitude at the inlet boundary, or the cruising speed, and *L*=0.25 m is the standard length of the flying fish. Three typical cases are chosen to present: (a) the rigid flapping tail at *f* = 145 Hz and *θ*_0_ = 10°, (b) the rigid flapping tail at *f* = 118 Hz and *θ*_0_ = 30° and (c) the periodically morphing tail at *f* = 98 Hz and *θ*_0_ = 30°. Blue (dark gray) and red (light gray) colors denote negative and positive pressure values respectively for 32 contour levels between −20000 and 5000 Pa.

We also present the drag forces in figure 3 (the dash-dotted lines), which are calculated by integrating the pressure and viscous forces along the flying fish excluding the contributions from its tail, or the propeller. For all cases, the drag forces are around −7.5 N. As suggested by Maertens et al. (2015) that it is very challenging to measure the efficiency for a self-propelled body in steady state, unless we can separate the propeller from the body. Here, assuming that the propulsive force produced by the tail is balanced by the drag force in steady state, we define the propulsive efficiency as

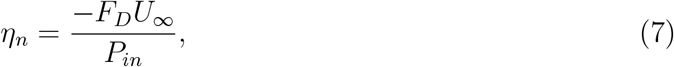

in which the power input *P*_*in*_ is obtained by integrating the inner product of distributed forces and moving velocities along the tail surface. For the steady states, or the cruising points identified in figure 3, we present their respective power input, or the power required to achieve the cruising states in figure 5. It is interesting to find that the required power increases with the maximum deflected angle, suggesting that the flying fish tends to consume less energy by beating its tail at a higher frequency rather than engaging a large amplitude. However, the minimum power input of 350 W shown in figure 5 requires a very high beating frequency of up to 145 Hz, which is far beyond the observed data (50 Hz), as well as that from the physiological constrain, as mentioned in section I. Nevertheless, from a hydrodynamic point of view alone, this minimum power is achievable. Actually, a real fish has more freedom of deformations than the model with a fixed incoming flow used in the current study, which could help reduce the required power when the deflected angle is large. Therefore, a self-propelled model might be more suitable (Deng and Caulfield 2018, 2016), which can be considered in future studies.

**FIG. 5:**
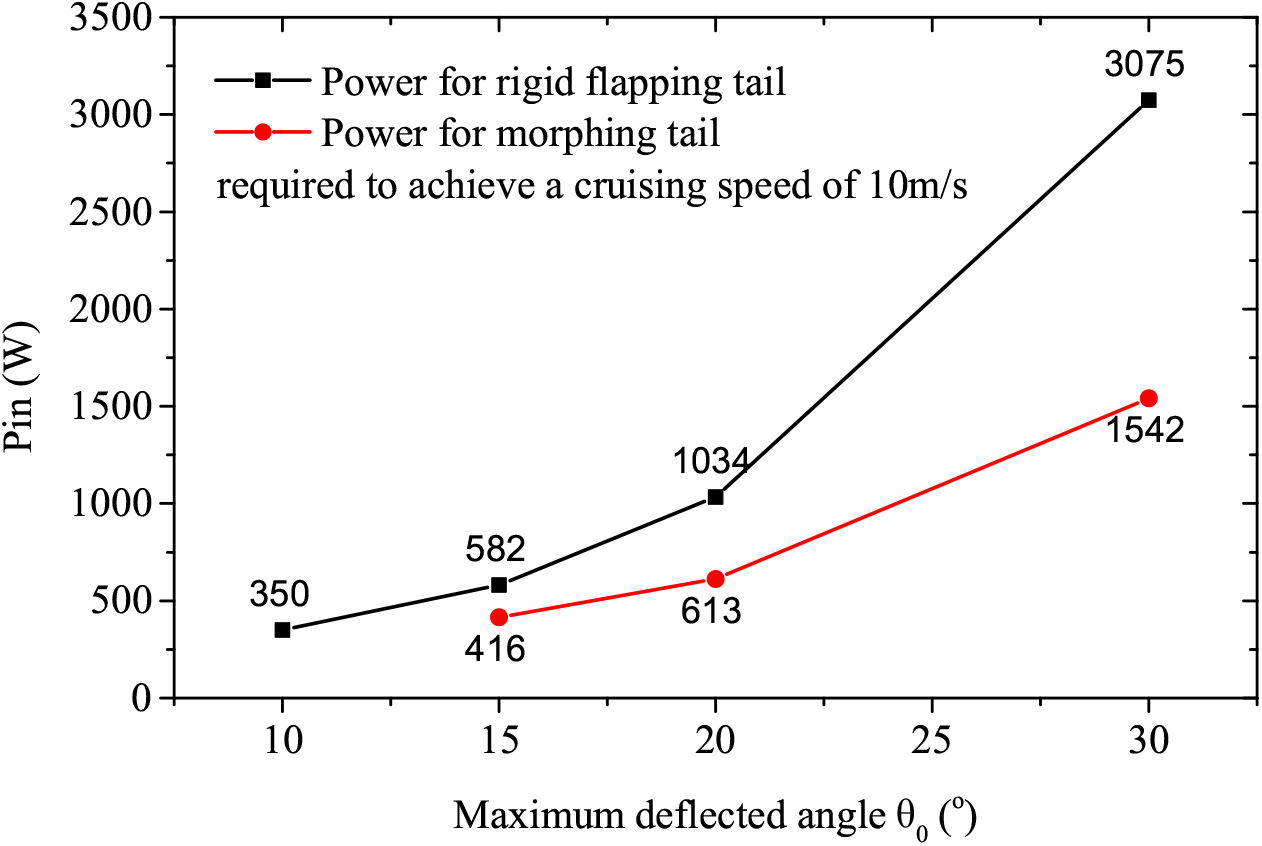
The power required for the flying fish swimming at a cruising (‘steady-state’) speed of 10 m s^−1^. Note that the cruising state is achieved when a zero horizontal force is obtained, i.e., the intersection points of the force curves and the dash lines shown in figure 3 (a) and (b).

### B. Surface taxiing locomotion

To understand the mechanism of further acceleration by taxiing on the water surface, here we study a flying fish beating its semi-submerged tail, with a fixed angle of attack *α* = 5° providing sufficient lift forces lifting up the fish from the water. The tail is bent down to an inclined angle of 30° with respect to the horizontal plane, as shown in figure 2. The tail performs periodically pitching motion, following the first propulsive mode employed by the underwater swimming locomotion. Intuitively, it is easy to understand that the flying fish will experience less resistance since most components, including the body and the pectoral fins, are airborne.

In figure 6, we show the variations of thrust forces with the beating frequency with two different *θ*_0_ considered. It is unsurprising to see that the critical frequencies for the flying fish to reach a cruising state, at the same speed of *U*_∞_ = 10 m s^−1^, are lower than that for the swimming locomotion (see figure 3 (a)). They are 50 Hz and 60 Hz for *θ*_0_ = 15° and *θ*_0_ = 10°, respectively. It seems that these frequencies accord more well with the observed data (50 Hz, as reported by Hertel (1966)).

**FIG. 6:**
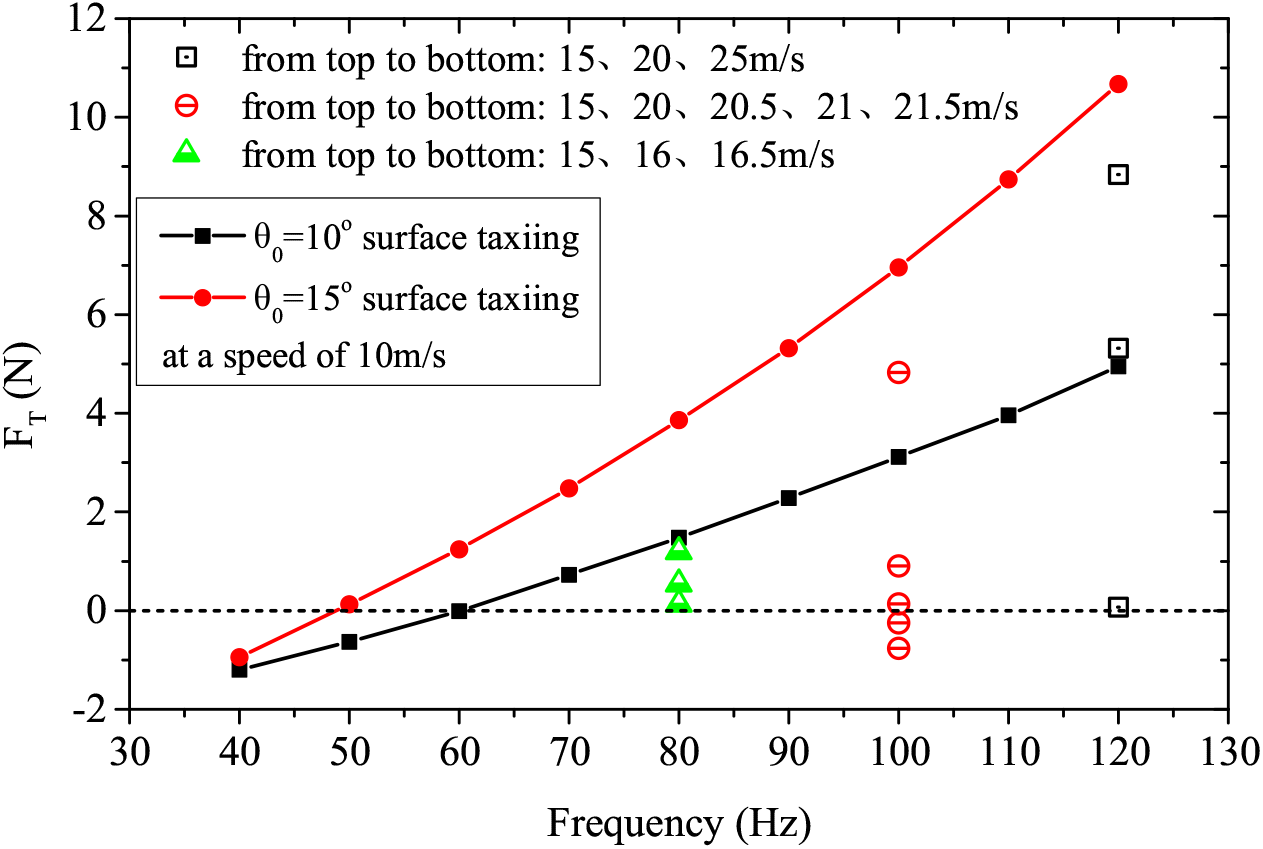
Horizontal forces on the flying fish when it taxis along the water surface for different beating frequencies. The lines marked with symbols show that for the flying fish taxiing at a speed of 10 m s^−1^, while the scattered symbols show that with various taxiing speeds aiming to identify the cruising states for three specific frequencies.

Furthermore, we fix the maximum deflected angle at *θ*_0_ = 15° and increase the flow incoming velocity, or equivalently the locomotive speed, at three typical frequencies of *f* = 80 Hz, 100 Hz and 120 Hz. It is clearly shown from figure 6 that the flying fish has the potential to cruise faster as the beating frequency is increased. For example, when the flying fish beats its tail at a frequency of 80 Hz, a cruising speed of 16.5 m s^−1^ can be achieved, and if the tail beats at a frequency of 120 Hz, the cruising speed can reach up to 25 m s^−1^, which is far beyond the existing data and out of the physical limit of the fish.

Another issue that we should note is the vertical force balance, i.e., between the lift force and gravity. For a specific case, at *θ*_0_ = 15°, *U*_∞_ = 16.5 m s^−1^ and *f* = 80 Hz, we find that the lift force provided by the pectoral fins is 2.08 N, or 0.21 kg in weight, which is very close to its body mass as mentioned in section 1. The drag force on the pectoral fins is 0.6 N, which is mainly induced by the lift, resulting in a lift-to-drag ratio of 3.47. Nondimensionlized by 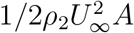, we find that the corresponding lift coefficient is 0.539, consistent with our previous study for a gliding flying fish at the same angle of attack of 5° (Deng et al. 2019). We thereby believe that the lift force is sufficient to lift up the flying fish undergoing a steady state taxiing. It is important to appreciate that the inclined tail fin can actually provide a small portion of lift force by propelling itself again the water, which is however not the major concern of the current study, and we leave it open to the future investigations.

In figure 7, we show the water-air interfaces for four typical taxiing cases, exhibiting unique wake patterns left on the glassy water surface, which have been widely observed by zoologists (Howell 2014). The clearly observed water splashes demonstrate sufficient resolution of our numerical method for the free surface.

**FIG. 7:**
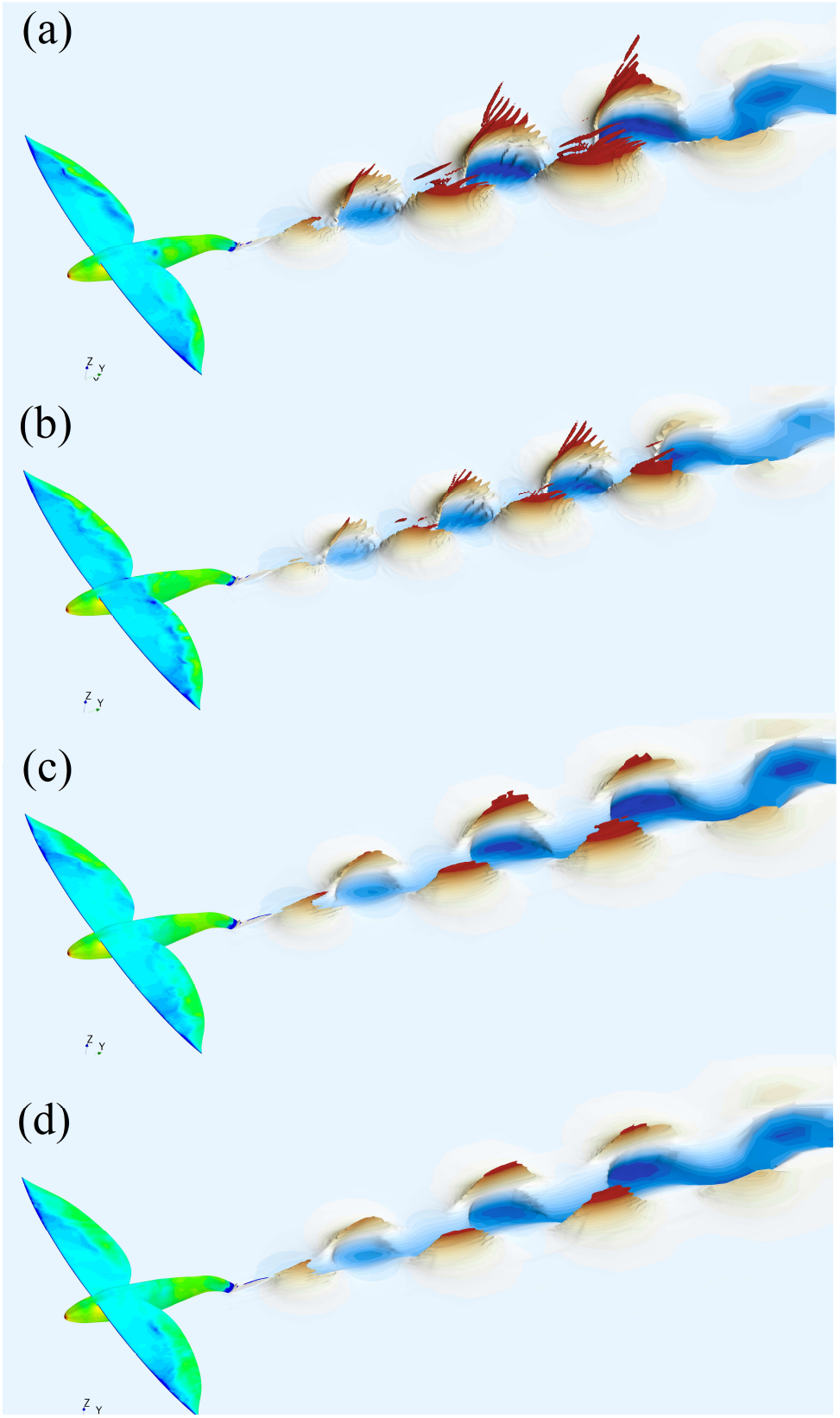
Water-air interfaces represented by the isosurface of volume fraction of water being equal to 0.5, with the colors from blue (dark gray) to red (light gray) denoting the depths varying from −0.05 m to 0.05 m. The different colors on the body and pectoral fins represent the pressure distributions. Four typical cases are shown. They are (a) *f* = 50 Hz, *U*_∞_ = 10m s^−1^, *θ*_0_ = 15°, (b) *f* = 60 Hz, *U*_∞_ = 10 m s^−1^, *θ*_0_ = 10°, (c) *f* = 80 Hz, *U*_∞_ = 16.5 m s^−1^, *θ*_0_ = 15°, and (d) *f* = 100 Hz, *U*_∞_ = 20.5 m s^−1^, *θ*_0_ = 15°.

In figure 8 we present the powers required to achieve respectively the five cruising points in figure 6. By making a direct comparison with figure 5, we see that the minimum power required to reach a taxiing speed of 10 m s^−1^ is 36 W, which is one order of magnitude smaller than that obtained for an underwater swimming flying fish. It is interesting to find that by inputting the same amount of power of 350 W, which is the minimum requirement for swimming locomotion, the taxiing fish is able to reach a cruising speed of up to 16.5 m s^−1^. Therefore, it is apparently evidenced that the flying fish can be further accelerated before take-off by taxiing on the water surface.

**FIG. 8:**
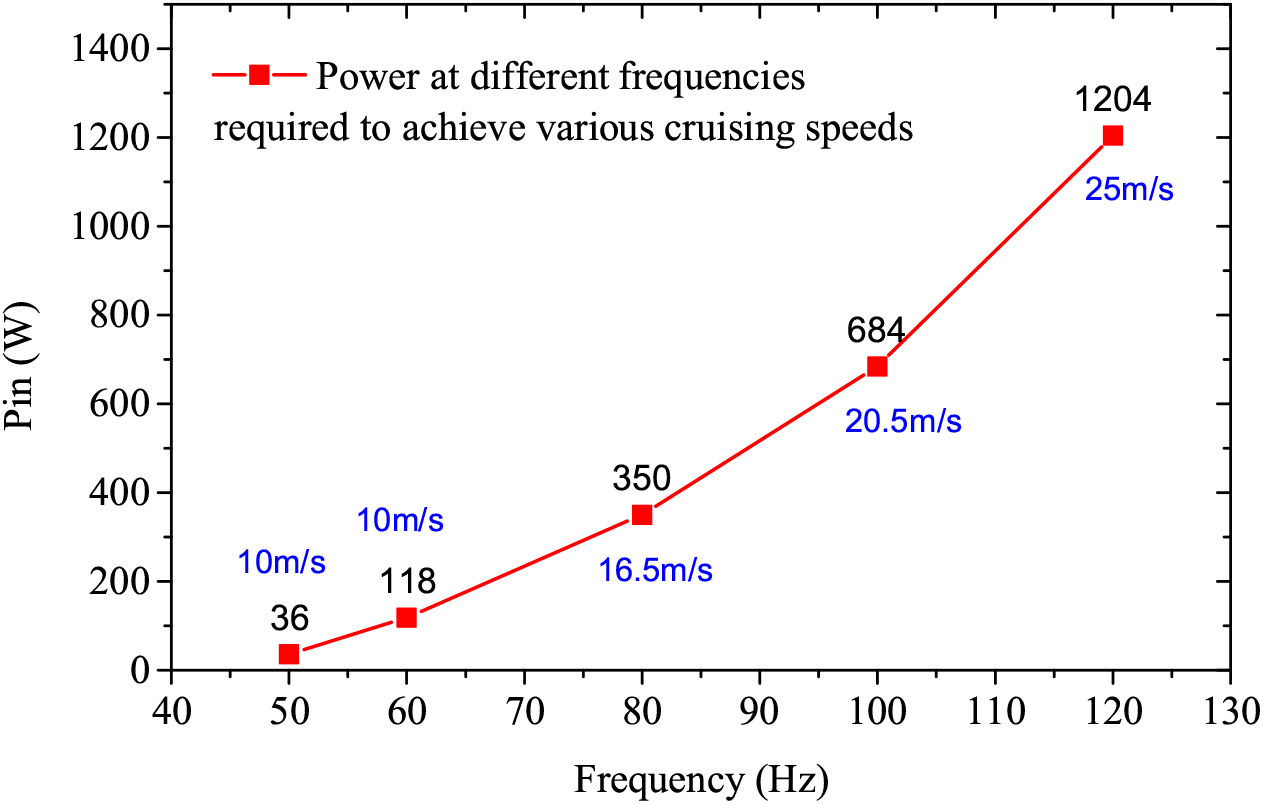
The power required for the flying fish taxiing at various cruising (‘steady-state’) speeds, corresponding to the five horizontal force balanced points in figure 6.

Assuming that the flying fish’s work capacity is constrained by its muscle power density, as we have mentioned above, and considering the currently studied flying fish with 0.191 kg in weight, the 350 W power relates to a muscle power density of 3664 W kg^−1^ (assuming 50% muscle by weight). Although this value is considerably higher than the muscle power density (2300 W kg^−1^) given by Gao and Techet (2011), we still believe that it is a reasonable estimation for a real flying fish at this length scale.

Moreover, we present the wake topologies for a typical case of surface taxiing in figure 9, exhibiting both the water surface distortion and the vortical structure of the flow. Besides the trailing edge shedding vortices from the spread pectoral fins, the finer vortex filaments induced by the interaction between the flow wake and the free surface are clearly observed.

**FIG. 9:**
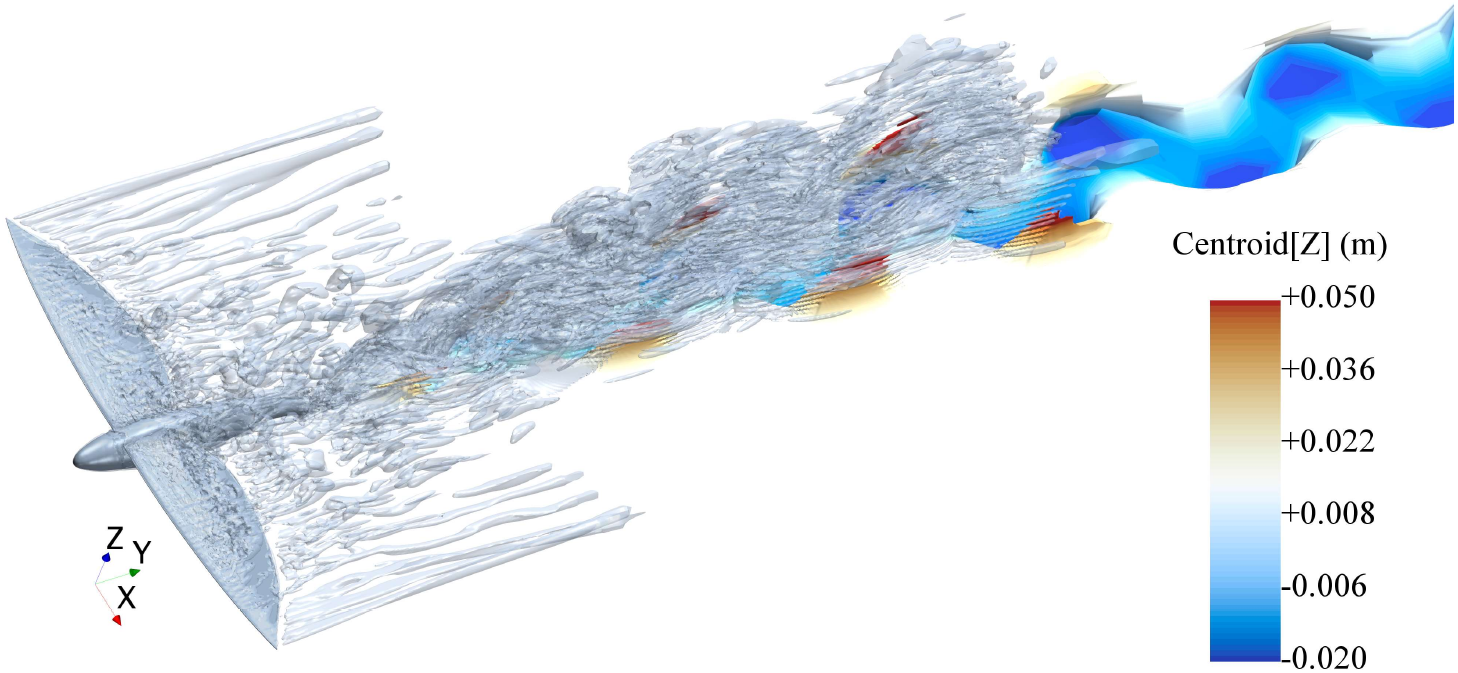
Wake topologies for the surface taxiing flying fish, visualized using *Q* = 7.5(*U*_∞_/*L*)^2^, in which *U*_∞_ =16.5 m s^−1^, and the flying fish beats its tail at the frequency *f* = 80 Hz. Blue (dark gray) and red (light gray) colors denote negative (below the still water surface) and positive (above the still water surface) depths respectively.

## IV. CONCLUSIONS

Following our previous study on the gliding performance of flying fish, here, we are concerned with their underwater swimming and water surface taxiing locomotion. Computa tional fluid dynamic (CFD) method is applied to investigate the hydrodynamic performance, focusing on its horizontal force balance and the power required to achieve a cruising state. We aim to answer the question ‘why does a flying fish taxi on sea surface before take-off’ from a hydrodynamic perspective.

First, for the underwater swimming locomotion, we consider two different propulsive modes and four different maximum deflected angles, at a constant speed of 10 m s^−1^. The results show that the critical frequencies are around 115 Hz for *θ*_0_ = 15°, 20° and 30°, while it is 145 Hz for the small deflected angle of *θ*_0_ = 10°, at which point the minimum power input of 350 W is recorded. We state that the required beating frequency, 145 Hz, for the minimum power is far beyond the data provided by field observation, which is around 50 Hz. Nevertheless, we suggest that it is achievable from a pure hydrodynamic point of view.

Second, we study a flying fish beating its semi-submerged tail fin, with a fixed angle of attack *α* = 5°, and a bent tail down to the water with an inclined angle of 30° with respect to the horizontal plane. It is unsurprising to find that the critical frequencies for *θ*_0_ = 15° and *θ*_0_ = 10° are 50 Hz and 60 Hz respectively, which are markedly lower than that of the swimming locomotion. Also they are much closer to the observed frequency of 50 Hz provided by previous zoologists. It sounds reasonable that the frequencies of surface taxiing are more likely to be recorded in field observation thanks to the wave patterns left on the glassy sea. It is exciting to find that by inputting the same amount of power of 350 W, which is the minimum requirement for swimming locomotion, the taxiing flying fish is able to reach a cruising speed of up to 16.5 m s^−1^. It is apparently evidenced that the flying fish can be further accelerated before take-off by taxiing on the water surface.

We understand that natural animals are unique with many unrevealed capabilities. It is a great challenge for artificial robots to wholly duplicate their locomotive mechanisms. The current study only sheds a first light to the aerial-aquatic locomotion of flying fish. We believe that there are more underlying physics lying behind flying fish waiting to be discovered.

## ACKNOWLEDGMENTS

This research has been supported by the National Natural Science Foundation of China (Grant No: 11772299).

